# Epigenetic reprogramming guides sexual dimorphism during floral development in *Silene latifolia*

**DOI:** 10.1101/2025.08.26.672086

**Authors:** Tomas Janicek, Vojtech Hudzieczek, Hana Polasek-Sedlackova, Marie Kratka, Vaclav Bacovsky

**Affiliations:** Department of Plant Developmental Genetics, Institute of Biophysics of the Czech Academy of Sciences, Kralovopolska 135, 612 65 Brno, Czech Republic; Department of Experimental Biology, Faculty of Science, Masaryk University, Kamenice 5, 625 00 Brno, Czech Republic; Institute of Biophysics of the Czech Academy of Sciences, Kralovopolska 135, 612 65 Brno, Czech Republic; National Centre for Biomolecular Research, Faculty of Science, Masaryk University, Kamenice 5, 625 00 Brno, Czech Republic; Institute of Experimental Botany of the Czech Academy of Sciences, Centre of Plant Structural and Functional Genomics, Slechtitelu 31, 779 00, Olomouc, Czech Republic

**Keywords:** histone modification, H3K4me1, H3K9me2, transcriptional activity, Pol-IIS2ph, white campion, *Silene latifolia*, dioecy, floral development, sexual dimorphism, high-content imaging, AI-image segmentation

## Abstract

Dioecy, the condition in which male and female individuals exist as separate plants, represents a fascinating and relatively rare reproductive strategy, offering unique opportunities to study the genetic and epigenetic regulation of sexual dimorphism. While sex determining genes underlying dioecy have already been described for several plant species, the role of epigenetic modifications in meristematic cell populations remains poorly understood. In this study we describe the spatio-temporal deposition of three epigenetic markers during early stages of floral development in model dioecious species *Silene latifolia*. We selected H3K4me1, H3K9me2 and active Ser2 phosphorylated form of RNA Polymerase II (Pol-IIS2ph), to assess levels of chromatin condensation and transcriptional activity of meristematic cells during key developmental stages. Utilizing the novel approach of an AI-assisted nuclei segmentation and high-content imaging we created a single-cell resolution atlas for male and female floral meristems. Our results show a relationship between transcription activity and sex determination during early meristem development. Moreover, our results suggest that H3K9me2 deposition in the developing meristem is linked to sex-specific chromatin reprogramming events, such as pollen mother cell formation during anther maturation. Overall, these results offer new insights into the role of chromatin regulation during floral meristem development and improves our understanding of sexual dimorphism in dioecious species.

## Introduction

Although relatively rare among flowering plants, dioecy represents a striking evolutionary strategy with profound implications for plant reproduction, genetic diversity, and sexual dimorphism (Renner 2014). Male and female individuals differ from each other in a broad spectrum of phenotypic traits, collectively known as sexual dimorphism, which includes variations in both primary and secondary sexual characteristics. (Punzalan and Hosken 2010). In plants, the primary sexual characteristics refer to reproductive structures - anthers and gynoecium, while the secondary sexual characteristics include morphological or physiological traits beyond the reproductive organs, including plant size, flower number, and other characteristics (Barrett and Hough 2013; Galfrascoli and Calviño 2020; Punzalan and Hosken 2010). Sexual dimorphism primarily results from sex-biased or sex-limited gene expression, which leads to differential trait development between males and females (Tanurdzic and Banks 2004). Genes with sex-biased expression often become enriched on sex chromosomes and can further contribute to sexually dimorphic traits, as demonstrated in *Drosophila melanogaster* (Innocenti and Morrow 2010) and mice (Yang et al. 2006). Sex-biased expression has been shown to vary across different tissues in animals (Naqvi et al. 2019), and has already been documented in several plant models, including *Asparagus* (Harkess and Leebens-Mack 2017; Harkess et al. 2015), *Coccinia grandis* (Devani et al. 2019), *Silene latifolia* (Prentout et al., 2023; Zemp et al., 2016) and brown algae *Ectocarpus* (Lipinska et al. 2015). Despite growing interest in sexual dimorphism in flowering plants, key aspects of the epigenetic architecture underlying the evolution of sex-biased expression remain unresolved (Hemenway and Gehring 2023).

The majority of sexually dimorphic traits in plants originate from sex-specific regulation of the stem cell population - shoot apical meristem (SAM). In plants, SAM is responsible for generating all above ground organs (leaf, stem and flower), and as such, is a continuously self-renewing cell niche. It is further divided into central, peripheral and rib zones. The central zone represents pluripotent stem cells and serves as a layer of progenitor cells that move into the peripheral zone, later differentiating into leaves or floral organs. The rib zone is responsible for the maintenance and growth of stem and vascular tissue (Steeves and Sussex 1989). Reproductive organs develop after the SAM transitions into the floral meristem (FM). The development of floral organs is accompanied by chromatin remodeling and progressive changes in gene expression (Cronk and Müller 2020; Hemenway and Gehring 2023). As a result, the general hermaphroditic flower is formed through the concentric arrangement of four floral organs: sepals, petals, stamens, and pistil (Coen and Meyerowitz 1991). In contrast, dioecious plants exhibit defects in reproductive structures or even complete loss of floral whorl(s) (Cronk and Müller 2020).

In *Silene latifolia*, a model organism for studying dioecy and sex chromosome evolution, sexual dimorphism in reproductive organs arises from sex-specific developmental arrests during flower formation. In males, suppression of the gynoecium results from arrested cell division in whorl 4, while in females, anther development is blocked at the early sporogenous stage (Farbos et al. 1997). The proposed model for sex determination mechanism assumes regulatory feedback loop between the X-linked *WUSCHEL*-like and the Y-linked *CLV3*-like genes (Kazama et al. 2022). This was later confirmed by two independent studies, offering deeper insight into the origin of dioecy and sex determination in *S. latifolia* (Akagi et al. 2025; Moraga et al. 2025). Sex-biased gene expression in *S. latifolia* has evolved for both autosomal and sex-linked genes, with most changes relative to hermaphrodites occurring in females (Zemp et al. 2016). This expression divergence likely reflects the resolution of intralocus sexual conflict during the evolution of dioecy. Comparative study of nine *Silene* species further suggests that male-biased expression emerges first, followed by female-biased expression as sexual dimorphism becomes established (Prentout et al. 2023).

Despite substantial progress in understanding of sex chromosomes evolution and sex-biased expression in *S. latifolia* (reviewed in Saunders and Muyle 2024), there is a missing link between molecular and morphological insights that would lead to precise characterization of developmental processes, allowing better understanding of epigenetic regulation in sex-determination and sex-biased expression, in space and time.

To address this, imaging-based approaches such as confocal microscopy in *Arabidopsis* have enabled precise quantification of chromatin and developmental changes during spore mother cell formation (She et al. 2013; She and Baroux 2015). These methods allow for high-resolution visualization of nuclear morphology, chromatin condensation, and the spatial dynamics of key epigenetic regulators. Recent advances in imaging platforms, such as light-sheet microscopy, and 3D reconstruction of a whole nuclei or cell, have further extended these capabilities to more complex and less tractable plant systems. In particular, semi-automated image segmentation together with faster image acquisition of confocal microscopy enabled large-scale, data-driven microscopic analysis of individual nuclei within heterogeneous tissues (Randall et al. 2022; Rutowicz et al. 2024). High-content imaging offers a higher level of automatization in both image acquisition and data analysis than more typical microscopy experiments described above (Way et al. 2023). While high-content imaging is routinely applied in human-focused research such as DNA replication and stability (Polasek-Sedlackova et al. 2022), in plants it has yet to firmly take root.

In this work, we analyze anatomical differences between sexes and chromatin-related changes during meristem-to-organ transition using high-content imaging microscopy with fully automated imaging and instance image segmentation. The combination of these three aspects enabled us a precise dissection of chromatin features such as transcription activity, assessing the level of open and close chromatin at single-cell resolution in whole mount tissue sections during 3 developmental stages for both sexes. We provide spatial-temporal analysis between individual flower organs related to chromatin organization. In particular, our objective is to address two developmental biology questions related to sex differences in *S. latifolia*: (i) do histone marks exhibit spatial sexual dimorphism during male and female flower development, and (ii) is developmental arrest in males and females maintained epigenetically through the subsequent development during organ differentiation? Besides, we show how high-content microscopy may assist in molecular studies concerning epigenetic regulation during plant development.

## Material and Methods

### Plant material

Adult male and female plants of *S. latifolia* (U17 inbred line) were used for all the tissue preparation steps. The U17 seeds were sterilized and germinated as described in (Bačovský et al. 2022), grown in a growth chamber with a 16 h light/8 h dark cycle at 22°C and collected after forming flower buds

### Sample preparation

Flower buds of *S. latifolia* were collected and immediately put into a cold 4% Formaldehyde fixation solution. After harvesting all samples, the vacuum was applied for one hour, or until all samples fully submerged into the formaldehyde solution. Next, the fixative was changed, and samples were kept in fresh formaldehyde at 4°C overnight. The next day, samples were washed in RNase free water and dehydrated by a series of washes with increasing concentrations of ethanol solutions (30, 50, 70, and 96%) each step for one hour. The samples were stained overnight with Eosin Y dissolved in 96% ethanol (0,1% Eosin Y final concentration). The next day, samples were washed 3 times in 2 hours in absolute ethanol (100%) and then transferred in solution with increasing ratio of Histoclear for one hour (25, 50, 75, 100%). All these steps were carried out on ice and the samples during washes and kept in a cold room. After the final Histoclear bath, the staples were removed from vials, transferred into cassettes for histology sectioning, and put into a 50:50 solution of molted paraffin and Histoclear for an hour. Samples were then moved into a pure molten paraffin and kept embedded overnight. The next day, the paraffin baths were changed two times and kept in the final paraffin bath overnight. The next day samples were moved into molds for sectioning. Samples were manually trimmed before sectioning into rectangle blocks. Sectioning was carried out on a manual microtome with a thickness set to 10 µm. Sections were laid onto a glass slide covered with water to ensure the proper morphology. The water evaporated on a hot plate at 40 °C. After drying, the slides were stored at 4 °C until deparaffinization. Deparaffination was carried out by two 10-minute baths in Histoclear and then re-hydrated in a reversed ethanol row into the final bath of 1% PBS solution (Fig. S1a).

### Immunolabeling

Slides were first permeabilized with a washing buffer (1xPBS, 1% Triton-X 100 1mM EDTA) for 1h. Prepared specimens were blocked in a 3% blocking solution (3% BSA (w/v) in 1xPBS, 0.3% Triton-X 100, 0.1mM EDTA) for 2h at RT and incubated with primary antibody diluted in 1% blocking solution (1% BSA (w/v) in 1xPBS, 0.1% Tween-20) for 72h. Primary antibodies against H3K4me1 (ab8895) and H3K9me2 (C15200154) were diluted in a ratio 1:200. The primary antibody against RNA Pol-IIS2ph (ab5095) was diluted in a ratio 1:500. After incubation in a primary antibody, slides were washed 3x10min in 1xPBS and incubated with the secondary antibody conjugated with Fluorescein (α-mouse;715-097-003) or Alexa 647 (α-rabbit; 111-606-003) diluted in 1% blocking solution 48h. After washing off the unbound secondary antibody residues 3x15 min in 1xPBs, slides were dehydrated in an ethanol series and mounted in Vectashield supplemented by 4′,6-diamidino-2-phenylindole (DAPI).

### Image acquisition and analysis

Images were acquired using ScanR inverted high-content screening microscope IX83 (Evident) equipped with wide-field optics, UPLXAPO dry objective (20x (NA 0.8), fast excitation and emission filter-wheel device for DAPI, FITC and Cy5 wavelengths, Lumencor Spectra X illumination system and digital monochrome ORCA FLASH (CMOS, Hamamatsu) camera with HW and SW autofocus. Images were captured in an automated fashion with the ScanR acquisition software (Evident, v3.4).

Following image acquisition, the automated image analysis was performed using ScanR analysis software (Evident, v3.4) with specifications described below. For intensity-based nuclei segmentation, the DAPI signal was used for the generation of an intensity-based mask to identify individual nuclei as main objects. This mask was then applied to analyze pixel intensities in different channels for each nucleus. For AI-assisted nuclei segmentation, the nuclei were segmented using a pre-trained neural network to separate difficult-to-distinguish nuclei within flower organs and meristem. The comparison between intensity- and AI-based segmentation can be found in Fig. S2. After segmentation, we performed automated multiparameter analysis within the ScanR analysis software to obtain fluorescent intensities of individual fluorescent channels. Afterwards, data were exported as a table and analyzed in Spotfire software (Tibco, v.10.5.0.72). Color-coded scatter diagrams were used to visualize differences between female and male parts. Diagrams were converted to Excel (Microsoft office 365) spreadsheets and statistically processed in R studio. For each flower stage, at least six individual meristem sections were used, with more than 1000 cells per replicate. Flowers of later stages were captured in multiple images with overlapping layers. The image stitching was processed in Adobe Photoshop (v26.9).

### Statistical Analysis

We used two-way ANOVA to measure the effect of sex and tissue type (and their interaction) on the signal intensity. Based on high-content imaging data and object counts, we calculated the effect size (Cohen’s *d*) to more accurately capture biologically meaningful differences between individual layers and organ primordia. This approach helps to account for the fact that statistical tests based solely on mean differences can produce highly significant *p*-values, even for small differences, when sample sizes are large. In such cases, statistical significance may not translate into a developmentally relevant effect. Effect size values are interpreted following (Cohen 2013): d = 0.2–0.3 indicates a small effect, 0.4–0.6 a medium effect, and ≥0.8 a large effect. In this context, positive d-values reflect higher fluorescence intensity in males (male bias), while negative values (–) indicate higher intensity in female meristems (female bias). Contrasts and effect-sizes (Cohen’s d) for pairwise comparisons between the sexes in each stage were calculated using R package.

## Results

We focused on three developmental stages (STG 3-4, STG 5-6, STG 7-8) during male and female development and assigned individual cell populations and flower organs based on the previously published data (Farbos et al. 1997; Grant et al. 1994). The accurate segmentation of nuclei is a prerequisite for quantifying chromatin states and transcriptional activity in complex plant tissues. When applying conventional threshold-based segmentation (TS) to *S. latifolia* floral meristems, we observed frequent failures to resolve densely packed or overlapping nuclei, particularly in developing anther primordia and the central regions of the floral meristem (Fig. 1a, left). These limitations resulted in underestimated nuclei counts and the generation of large, unresolved clusters of signals (Fig. 1b, insets).

**Figure 1.**
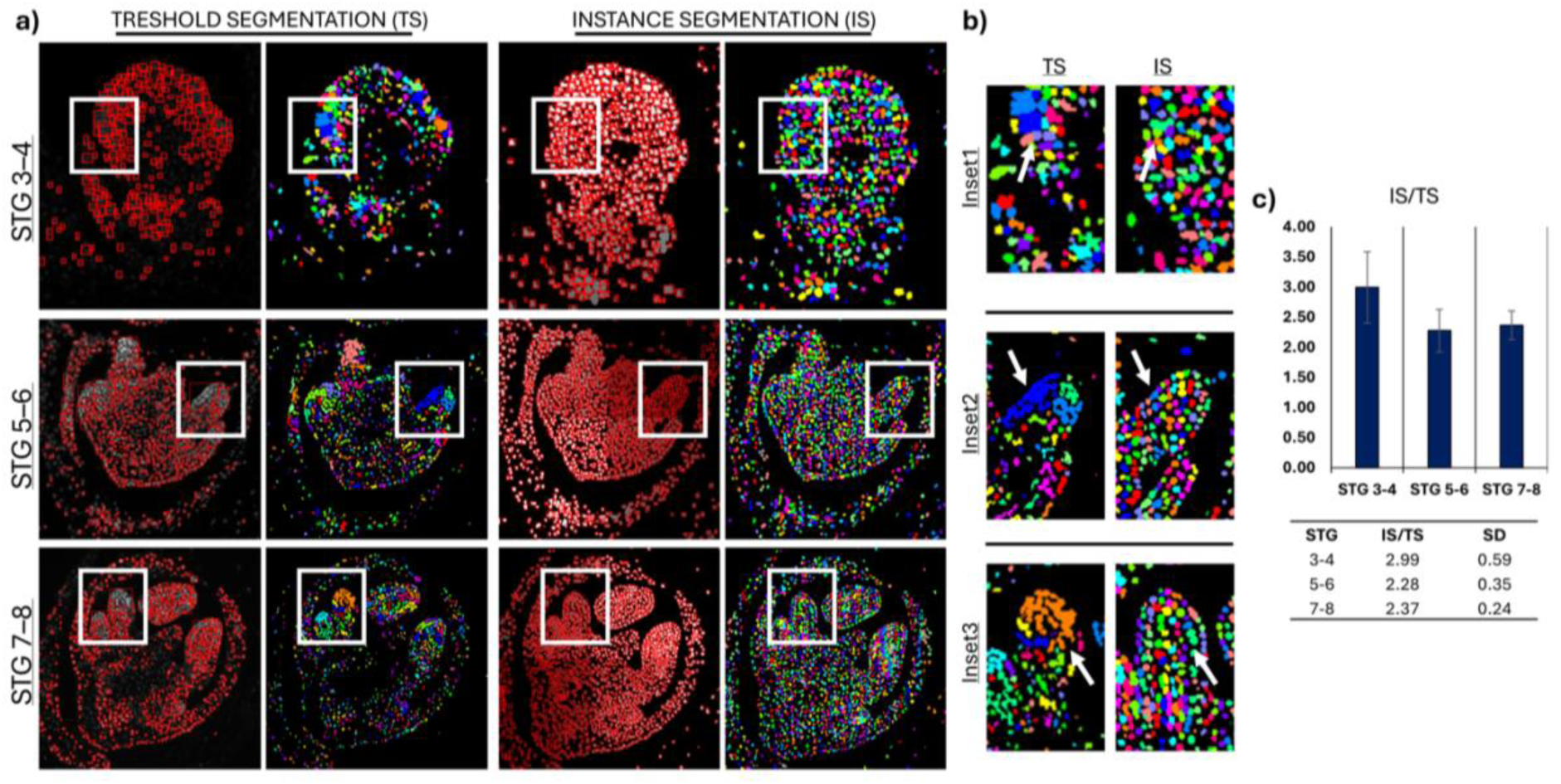
Comparison of threshold and AI-assisted instance segmentation for nuclei detection in S. latifolia floral meristems. (a) Representative images of floral meristems at stages STG 3–4, STG 5–6, and STG 7– 8, showing comparison between conventional threshold-based segmentation (TS, left) and TruAI-assisted instance segmentation (IS, right). Pol-IIS2ph and DAPI intensity is shown with a common scale. TS frequently underestimates nuclei counts in dense regions and merges overlapping objects, whereas IS resolves individual nuclei across all stages. (b) Insets highlight typical segmentation errors with TS (arrows), such as unresolved clusters or merged objects, contrasted with accurate separation achieved by IS. (c) Quantitative comparison of detected nuclei shows a consistent 2–3-fold increase with IS compared to TS, confirming the improved accuracy and robustness of AI-assisted segmentation.

To address these shortcomings, we implemented TruAI-assisted instance segmentation (IS). In contrast to thresholding, IS successfully separated touching and overlapping nuclei, providing more accurate object boundaries across all examined developmental stages (STG 3–4, 5–6, and 7– 8) (Fig. 1a, right). Detailed inspection of individual regions confirmed that TruAI resolves complex objects that remained unresolved with TS (Fig. 1b, arrows). Quantitative comparison between the two methods revealed a consistent 2–3-fold increase in detected nuclei with IS relative to TS (Fig. 1c), underscoring the robustness of AI-assisted segmentation.

For the respective stages, we compared the level of available active subunits of Pol-IIS2ph (Pol-II) and two histone modifications that are linked to active (H3K4me1) and repressed chromatin (H3K9me2), respectively. Among the tested stages (Fig. 2, 3, 4) and individual cell lines, we selected sepal tissue for data normalization due to the following characteristics - (i) sepals are present and relatively easy distinguishable in all studied stages, (ii) sepals displayed the most consistent results comparable between replicates and sexes, and (iii) sepals are comparable within other tissues that are present either only in males or females.

**Figure 2.**
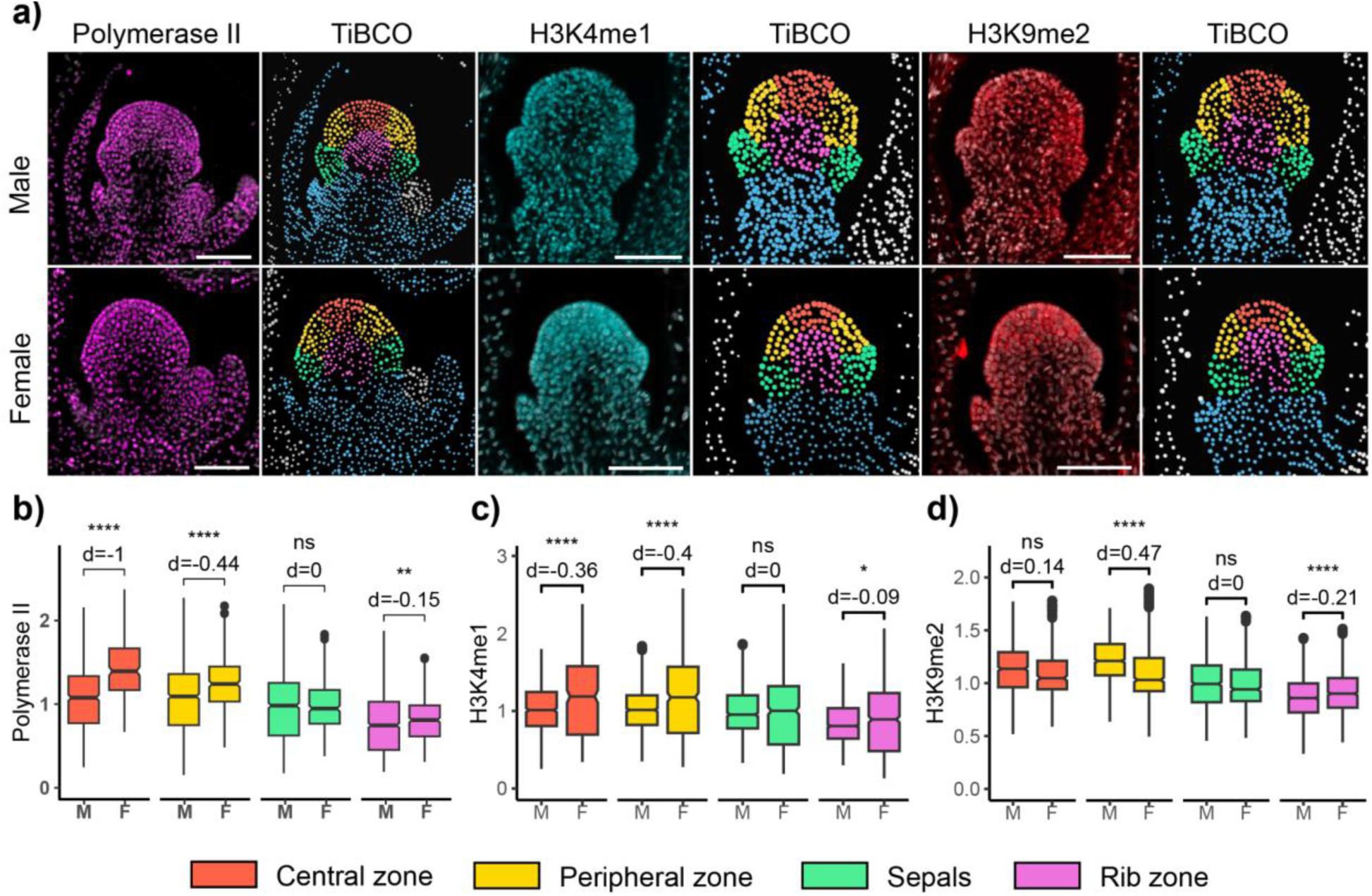
Longitudinal section of *S. latifolia* floral meristems at developmental stages 3–4. (a) Differences between male and female floral meristems in the distribution of Pol-IIS2ph (magenta), H3K4me1 (cyan), and H3K9me2 (red). Wide-field microscopic images were transformed into 2D scatter plots to visualize fluorescence signal intensities across distinct meristematic zones. (b–d) Box plots showing fluorescence intensity of each marker across individual meristematic layers. Individual cells were semi-automatically assigned to the central zone (red), peripheral zone (yellow), sepals (green), and rib zone (purple) in TIBCO Spotfire, with colors corresponding to those shown in (a). The distribution patterns of Pol-IIS2ph, H3K4me1, and H3K9me2 were consistent across biological replicates and individual meristems.

**Figure 3.**
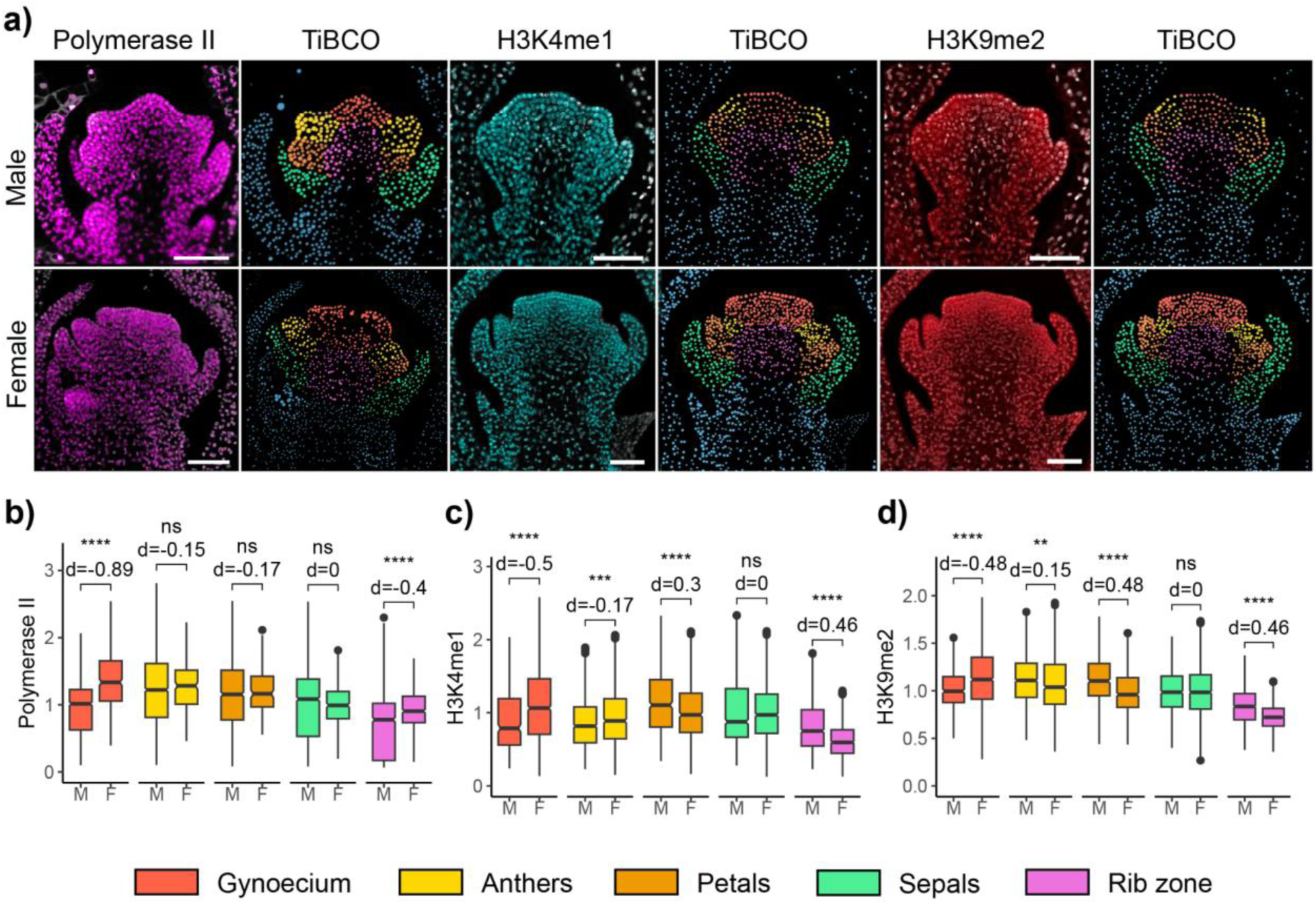
Longitudinal section of *S. latifolia* floral organ primordia at developmental stages 5–6. (a) Differences between male and female floral meristems in the distribution of Pol-IIS2ph (magenta), H3K4me1 (cyan), and H3K9me2 (red). Wide-field microscopic images were transformed into 2D scatter plots to visualize fluorescence signal intensities across distinct flower organs. (b–d) Box plots showing fluorescence intensity of individual fluorescent markers across individual meristematic layers. Individual cells were semi-automatically assigned to the gynoecium (red), anthers (yellow), petals (orange), sepals (green), and rib zone (purple) in TIBCO Spotfire, with colors corresponding to those shown in (a). The distribution patterns of Pol-IIS2phh, H3K4me1, and H3K9me2 were consistent across biological replicates and individual meristems. Note the differences between gynoecium flower organs in males and females for Pol-II and the local distribution of H3K4me1 in anthers.

**Figure 4.**
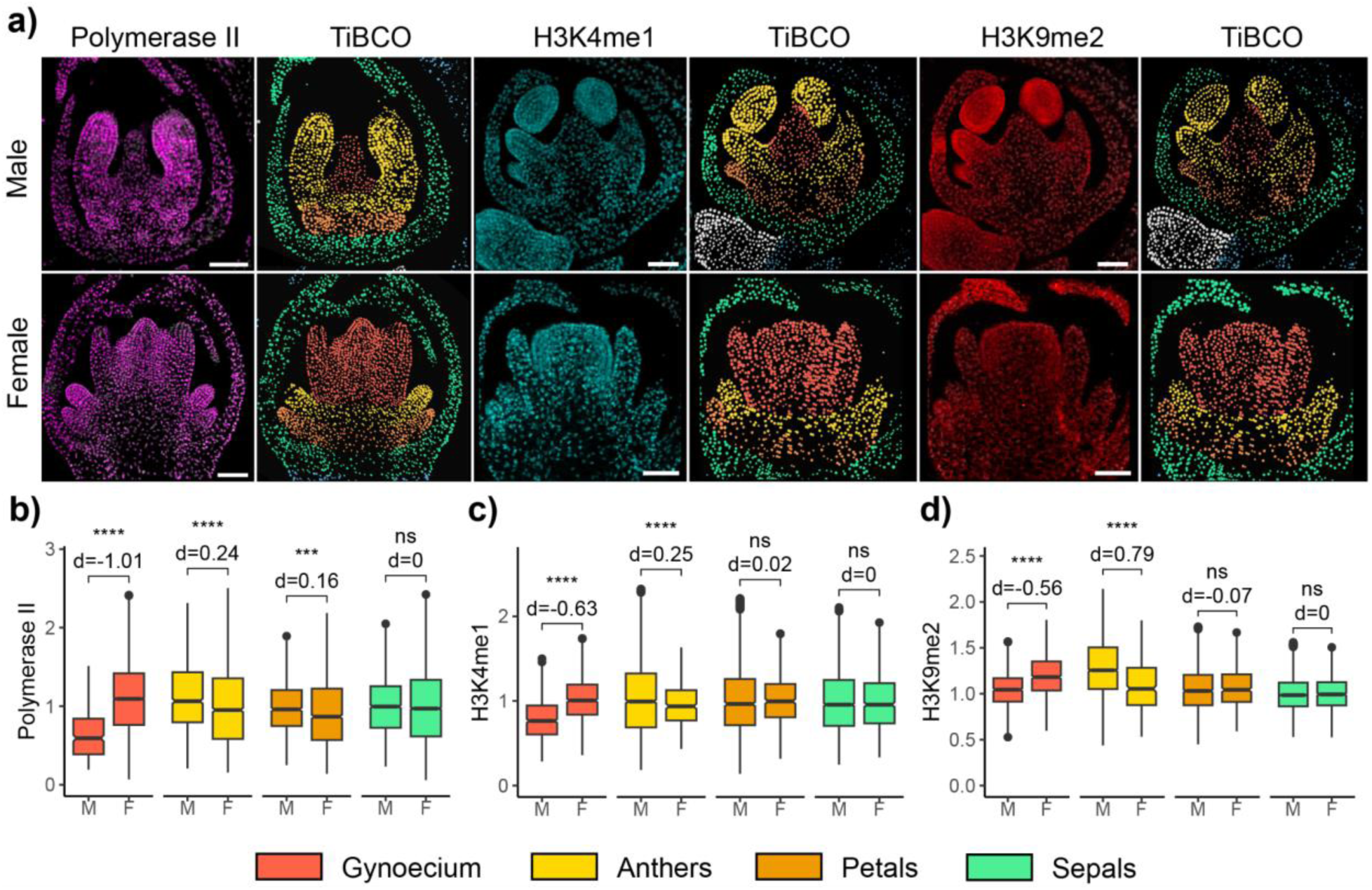
Longitudinal section of *S. latifolia* floral meristems at developmental stages 7–8. (a) Differences between male and female floral meristems in the distribution of Pol-IIS2ph (magenta), H3K4me1 (cyan), and H3K9me2 (red). Wide-field microscopic images were transformed into 2D scatter plots to visualize fluorescence signal intensities across distinct flower organs. (b–d) Box plots showing fluorescence intensity of individual fluorescent markers across individual meristematic layers. Individual cells were semi-automatically assigned to the gynoecium (red), anthers (yellow), petals (orange) and sepals (green) in TIBCO Spotfire, with colors corresponding to those shown in (a). The distribution patterns of Pol-IIS2ph, H3K4me1, and H3K9me2 were consistent across biological replicates and individual meristems. Note the differences in Pol2 distribution between gynoecium flower organ in males and females, and aborted growth of anthers with overall lesser abundance of H3K4me1 in females. Interestingly, the anthers in males are enriched for H3K4me1 and H3K9me2.

### Enhanced transcriptional activity and open chromatin state in female organ primordia support early development

Floral development in *S*. *latifolia* begins during developmental stages 3–4 (STG 3–4) (Fig. 2a). At this stage, male and female meristems are morphologically indistinguishable (Fig. 2a), with no visible signs of sexual dimorphism. At this stage, sepals are initiated as the first flower organs. The rest of the floral meristem (FM) is still comparable to the prior vegetative structures of SAM. Thus we separated cells into four groups utilizing sepals organ primordia with a central, peripheral and rib zone, characteristic cell groups within the SAM (Fig. 2a) (Steeves and Sussex 1989).

Based on the male to female (M:F) comparison from obtained data, we observed a statistically significant enrichment of active Pol-IIS2ph subunits in the female meristem, particularly in the central zone (*p* = 0.001) with a large effect size (*d* = -1.0) (Fig. 2b). The peripheral zone (*p* = 0.001) and rib zone (*p* = 0.15) also showed statistically significant differences, observable in correlation between DAPI and Pol-II mean intensity (Fig. S2). However, the effect sizes were medium only for the peripheral zone (*d* = -0.44) and weak for the rib (*d* = -0.15). No statistically significant differences were detected in the sepal region between sexes.

Based on the observed Pol-II enrichment in the central zone, we expected a corresponding increase in H3K4me1, a histone modification associated with open chromatin, in the same region of the female meristem (Fig. 2c). Although the differences between sexes were statistically significant (*p* = 0.001), the effect size reached a medium level only in the peripheral zone (*d* = -0.4). Interestingly, this lower enrichment of H3K4me1 in the female peripheral zone was accompanied by increased levels of H3K9me2, consistent with male-biased enrichment and a medium effect size (*p* = 0.001, *d* = 0.47). Correlative analysis between both histone marks visualized in 2D scatter plots support this trend (Fig. S3). In contrast, the enrichment of H3K9me2 in the central and rib zones showed very small effect sizes (*d* = 0.14 and *d* = -0.14, respectively). These results indicate that the central zone of the female meristem at STG 3–4 sustains high transcriptional activity (Fig. 2b, *d* = -1.0; Fig. S2, S3), along with elevated H3K4me1 levels, suggesting a more open chromatin state compared to males. Conversely, the peripheral zone in males appears to exhibit a more condensed chromatin state (Fig. 2d, Fig. S3), while females show low-to-medium enrichment of both Pol-II subunits and H3K4me1 in this region (Fig. 2b, c).

### Chromatin reorganization underlies male organ development at STG 5–6

The development of male and female floral organs at stages 5–6 (STG 5–6) is marked by the formation and co-occurrence of all remaining floral organ primordia (Fig. 3a). During these stages, the female floral meristem becomes noticeably larger than the male counterpart. At stage 5, petals and anthers begin to form, with the anthers originating from the peripheral zone established at stage 4. The most prominent difference in floral meristem organization lies in the size of the fourth whorl, which gives rise to the carpel primordia. By stage 6, the carpel primordia are already fully discernible in female meristems of *S. latifolia*. In contrast, the male meristem develops a modified, stigma-like filamentous structure in place of a carpels, reflecting its suppression. The emergence of anthers from peripheral cells and the absence of a gynoecium in males represent two sequentially expressed dimorphic traits, linked to differences in meristem size and organization. At STG 5–6, we classified cells according to their floral organ primordium identity, including the rib zone at the center of the floral meristem (Fig. 3a).

During STG 5–6, the transcriptional activity and chromatin state in individual organ primordia largely follow the patterns observed at STG 3–4 (Fig. 3a–d), with additional chromatin-related suppression associated with gynoecium development in females (Fig. 3a, d). The M:F enrichment of Pol-II subunits shows a statistically significant bias toward the gynoecium and rib zone (*p* = 0.001), with a strong effect size in the gynoecium (*d* = -0.89) and a medium effect in the rib zone (*d* = -0.4). Global distribution of Pol-II and the relationship between DAPI mean intensity support this trend, with lower intensity for both organs, suggesting a more relaxed chromatin state (Fig. S4). This transcriptional activity is supported by H3K4me1 enrichment (*p* = 0.001, *d* = -0.5), although signs of chromatin reorganization are apparent in the rib zone of males (Fig. S4, S5). The M:F comparison for H3K4me1 in the rib zone also shows a significant difference (*p* = 0.001) with a medium effect (*d* = 0.46).

H3K9me2 levels increase in the gynoecium primordia of females (*p* = 0.001, *d* = -0.48) and in the male rib zone (*p* = 0.001, *d* = 0.46). Anthers and petals show low or no statistical significance and weak effects (Fig. 3b, c), except for a general increase of H3K9me2 in male petals (*p* = 0.001, *d* = 0.48; Fig. 3d). These results suggest that the female gynoecium remains transcriptionally active while undergoing chromatin reorganization, marked by increased global levels H3K9me2 (Fig. 3d, Fig. S5). Although transcriptional activity and open chromatin in anthers are not significantly different between sexes (*d* = -0.15 for both Pol-II and H3K4me1), there is a minor trend toward increasing global transcription in male organs (Fig. S4).

### Stage 7–8 reveals divergent chromatin and transcriptional dynamics in male and female floral organs

The final group, comprising stages 7 and 8 (STG 7–8), is characterized by continued growth of floral organ primordia. This phase marks the transition to reproductive development, with the onset of pollen and ovule formation. In male flowers, anther lobes begin to differentiate, while the arrested gynoecium elongates into a central filamentous structure resembling a stigma. In contrast, female flowers exhibit continued gynoecium enlargement, accompanied by a gradual suppression of anther development.

Transcriptional activity and open chromatin state remain significantly biased toward gynoecium development at STG 7–8 in females *(p =* 0.001), with a strong effect size *(d =* -1.01) (Fig. 4a, b). The same holds at the global levels of Pol-II in relation to DAPI, namely for gynoecium and petals (Fig. S6). This is accompanied by enrichment of H3K4me1 *(p =* 0.001*, d =* -0.63). As expected, gynoecium development in males is aborted (Fig. 4a), as visualized by the reduced fluorescence signal in the TiBCO scatter plot (Fig. 4a), and biased nuclear levels toward repressed chromatin (H3K9me2; Fig. S7). Although the M:F effect size for Pol-II abundance in anther primordia is weak, it begins to shift in favor of male development *(p =* 0.001*, d =* 0.24) (Fig. 4b; Fig. S6). Compared to STG 5–6, anther primordia at STG 7–8 show increased H3K4me1 levels in males *(p =* 0.001*, d =* 0.25), supported in comparison of both histone marks and global increase of H3K4me1 (Fig. S7).

Interestingly, H3K9me2 levels increase in the female gynoecium *(p =* 0.001*, d =* -0.56) and in male anthers *(p =* 0.001*, d =* 0.79). No significant differences are observed in petals and sepals (Fig. 4b–d). Given the strong fluorescence signal maxima in male anthers for both Pol-II and H3K4me1, compared to the strongly reduced female gynoecium, STG 7–8 exhibits the most pronounced M:F differences in chromatin organization and global transcription (Fig. 4, S6, S7). Although the overall effect sizes for active transcription markers are lower at STG 7–8 compared to STG 3–4 and STG 5–6, both Pol-II and H3K4me1 remain biased toward male organs. The global transcriptional activity in the female gynoecium is not surprising, given the fact that male gynoecium development is largely aborted (Fig. 3a, 4a). Notably, the strongest enrichment of Pol-IIS2ph and H3K4me1 is observed in the central region of the female gynoecium, specifically in the area corresponding to the future placenta and ovules (Fig. 4a).

When comparing the global effect sizes for M:F differences across developmental stages and fluorescent markers, the most pronounced differences are observed for Pol-II and H3K9me2. Pol-II is consistently biased toward female organ primordia, with a substantial decrease in signal intensity and a shift toward male-biased expression by STG 7–8 (Fig. 5). A similar but inverse pattern is seen for H3K9me2, which is biased toward male organ primordia, though its effect becomes more neutral at STG 7–8. While H3K4me1 generally shows a bias toward female organs, the pattern becomes more organ-specific, with shifts in bias toward male development at later stages (Fig. 3c, 4c). These epigenetic shifts coincide with key morphogenetic events, for instance, the Pol-II and H3K4me1 enrichment in the gynoecium primordia supports early ovule and placenta initiation in females at STG 7–8. In contrast, the later H3K9me2 accumulation in anthers may reflect chromatin compaction linked to pollen sac differentiation in males. Thus, sexually dimorphic chromatin landscapes not only reflect but likely reinforce divergent developmental programs, ensuring proper formation of sex-specific reproductive structures, from early meristematic stage (Fig. 2) to meristem-to-organ transition stage (Fig. 3), and organ primordia differentiation (Fig. 4).

**Figure 5.**
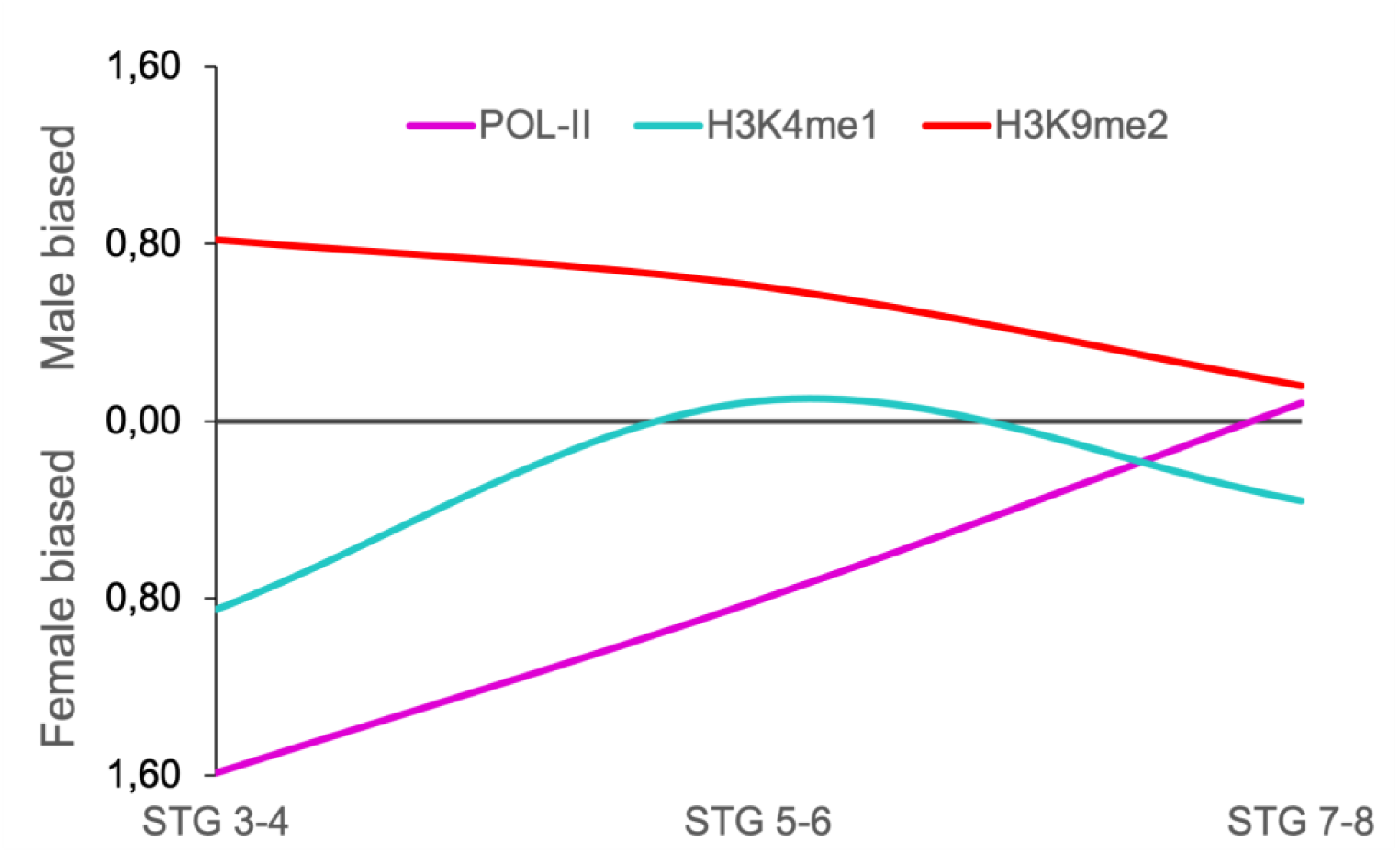
Comparison of total effect size between male and female flowers across developmental stages. The figure illustrates sex-specific biases in the abundance of Pol-II and individual chromatin markers and their dynamics over time. During the meristem-to-organ transition in females (stages 3–4) and early organ differentiation (stages 5–6), female meristems and developing organs show a bias toward higher levels of active Pol-IIS2ph subunits. In later stages, corresponding to intensified growth in males, this bias shifts toward male organs, accompanied by an increase in H3K4me1 levels and a decrease in H3K9me2.

## Discussion

Sexual dimorphic traits occur in plants throughout the development of the individual as the reproductive organs emerge and differentiate. Studies on *Drosophila* and other model species revealed the sex-biased expression changes from one developmental stage to another (Grath and Parsch 2016; Naqvi et al. 2019; Zhou and Bachtrog 2012). However, it is recognized that further studies on relation of expression data, fitness and development are needed to resolve the evolutionary significance of sexual biases (reviewed in Saunders and Muyle 2024). The limited knowledge of the regulatory networks underlying sexual dimorphism during floral development makes further conclusions on the relation of sex-specific fitness and expression rather difficult. As a well-established model for plant sex chromosome biology, *Silene latifolia* has greatly contributed to our understanding of sex-specific gene expression (Zemp et al. 2016), dosage compensation (Muyle et al. 2018), and the evolution of sex determination (Kazama et al. 2022). Building on this foundation, we investigated epigenetic variation by quantifying global levels of Pol-IIS2ph and two chromatin-associated histone marks, H3K4me1 (open) and H3K9me2 (repressed), in the young sex chromosome system of *S. latifolia*.

### Expression and active chromatin levels are biased towards early organ primordia development in females

Floral development in *S*. *latifolia* has been previously divided into twelve distinct developmental stages (Farbos et al. 1997; Grant et al. 1994; Zluvova et al. 2006). While each morphologically unique, early floral stages often share developmental programs. Therefore, we grouped the stages into three broader categories likely to undergo similar epigenetic regulation, displaying similar transcriptional profiles: stages 3–4 (STG 3–4), 5–6 (STG 5–6) and 7–8 (STG 7–8). These represent the undifferentiated meristem stage (STG 3–4), the transition from meristem-to-organ primordia and the onset of sex-specific developmental arrest (STG 5–6), and the period of male and female organ differentiation (Farbos et al., 1997). The organ differentiation includes the development and enlargement of anthers and connective in males, and the closure of the ovary and the occurrence of stigma and differentiated placenta (STG 7–8).

At STG 3–4, no morphological differences are visible between male and female floral meristems (FM) (Fig. 2), corresponding to previously described characteristics (Farbos et al. 1997). However, gene expression and chromatin state analysis in *Arabidopsis* has shown that even SAM and early-stage meristems can exhibit sex-specific expression patterns and chromatin remodeling (reviewed in Hemenway and Gehring 2023), particularly as organ primordia begin to expand. In *S. latifolia*, female meristems at STG 3–4 exhibit elevated levels of the Pol-II Ser2-phosphorylated subunit (Pol-IIS2ph) in the central zone, while male meristems show comparatively lower transcriptional activity (Fig. 3b). This transcriptional bias in females is supported by more accessible chromatin states, as indicated by high H3K4me1 levels (Fig. 2c, Fig. 5). At the same time, female meristems display low to moderate H3K9me2 enrichment, in contrast to higher levels observed in male tissue (Fig. 2d). These differences are most pronounced in the central and peripheral zones, with both Pol-IIS2ph and H3K4me1 consistently biased toward female development (Fig. 5).

The higher expression activity and open chromatin state align with the expression of previously characterized genes known to govern FM maintenance and organ differentiation. These include *S. latifolia WUSCHEL* (*SlWUS1*), *SHOOT MERISTEMLESS 1* and *2* (*SlSTM1*, *2*), and *SLM2* gene, a homologue to the B-class MADS-box gene *PISTILLATA* (reviewed in Hobza et al. 2018). While the central zone in males lacks the expression of *SlSTM1*, females and hermaphrodites sustain its expression till the end of stage 4 (Zluvova et al. 2006). Interestingly, *SLM2* shows sex-specific differences in location and intensity, being enriched in the central part of the peripheral zone in females while in males it borders the central zone with much higher expression levels, based on in situ hybridization patterns (Hardenack et al. 1994). These spatial expression differences may reflect early regulatory events driving floral organ identity. We therefore propose that the increased levels of Pol-IIS2ph and H3K4me1 are linked to the proliferative phase prior to floral organ initiation, while the H3K9me2 serves as a marker for stem cell determination (Finnegan et al. 1996; Loh et al. 2007). This would point to an active role of these markers in the aforementioned transcriptional networks.

The developmental stage STG 5–6 marks the emergence of clear sexual dimorphism during meristem-to-organ transition. In these two stages, the development of gynoecium in males is clearly arrested, consistent with earlier reports (Farbos et al. 1997; Grant et al. 1994). We observed the same developmental arrest, accompanied by the increased levels of H3K9me2 in the rib zone, with a moderate male-biased effect size (*d =* 0.46) (Fig. 3d). The observed transcriptional activity trend aligns with the floral expression and continued expansion of the gynoecium in female flowers, which is expected to be highly transcriptionally active at this stage (Berger and Twell 2011). However, the repressive marker H3K9me2 showed medium increase in female 4th whorl as well. This could be the result of stem cell determination, as the gynoecium is the last primordia to be set in *S. latifolia* (Donnison et al. 1996). The second possible cause for the increased H3K9me2 could be the onset of the *AGAMOUS* (*AG*) expression which has been previously linked to the higher methylation levels in respective tissue (Finnegan et al. 1996; Hardenack et al. 1994; Jacobsen et al. 2000). This process could mark the beginning of the pre-meiotic developmental program, ultimately leading to the speciation of spore mother cell (SMC) in male or megaspore mother cell (MMC) in the female germline (She and Baroux 2014). Since the repression of somatic program is established de novo within reproductive cell niche and future floral sex organs, MMC differentiation is accompanied with increase of H3K9me2, in addition to H3K4me3 enrichment and the loss of H3K27me3 before the cells enter meiosis (She et al. 2013; She and Baroux 2014) 2014). In *Arabidopsis*, H3K9me2 is maintained through the action of histone-lysine N-methyltransferase *SUVH4*. Interestingly, the *Silene* ortholog of *SUVH4* was found to be upregulated in later female developmental stages (STG9+) (Bačovský et al. 2022). A moderate decrease in expression of this gene in *S. latifolia* androhermaphrodites supports a conserved role for H3K9me2 in female organ development aligning with other plant species (Berr et al. 2011). Both petals and sepals did not exhibit major differences between sexes, although H3K9me2 and, to a lesser extent, H3K4me1 levels were moderately elevated in male petals, with small to medium size effect (Fig. 3c, d).

### Sex specific differences during organ development

The end of stage 8 is characterized by formation of specialized meiotic cells that further undergo rapid chromatin reorganization distinct from their somatic progenitors (Farbos et al. 1997; Lloyd and Lister 2022). A similar pattern is evident in female *S. latifolia* FM at STG 5-6 during gynoecium primordia formation, and in male anthers at STG 7–8, which are likely entering meiotic development (Ashapkin et al. 2019; Loh et al. 2007). During STG 7–8, organs in both sexes continue to expand. The most prominent sexually dimorphic feature is the suppression of anthers development in female floral meristem (Fig. 4a), corroborating again previous findings (Farbos et al. 1997).

The levels of active subunits of Pol-IIS2ph and both histone modification intensities are higher in female organ primordia, with moderate decrease compared to the previous stages. The Pol-IIS2ph becomes increasingly localized to the central region of future placenta and ovary (Fig. 3b). In contrast, in male flowers, we expected higher Pol-IIS2ph abundance in the 3rd whorl corresponding to anther development. Interestingly, the most pronounced chromatin difference was a decrease in H3K9me2 levels, accompanied by weak-to-moderate increase of H3K4me1, both biased towards developing anthers (Fig. 4c, d; Fig. 5). These bimodal epigenetic patterns are consistent with previously reported epigenetic transitions during gametogenesis, where a reduction in repressive histone marks has been documented in both sexes, or more prominently in females, in cells that give rise to ovules and surrounding nutritive tissue similarly as in this study (Fig. 3). In *Arabidopsis*, both the male and female gametophytes undergo rapid chromatin and transcriptional reprogramming prior to meiosis. The typical characteristics are marked by the erasure of repressive histone marks (H3K27me1, H3K27me3 and H3K9me2) favoring active transcription, together with the enrichment of H3K4me2 and H3K9ac (Lloyd and Lister 2022; She et al. 2013). Therefore, we hypothesize that the observed sex-biased levels in H3K9me2 and H3K4me at STG 7–8 reflect the onset of gametophytic developmental programs, namely microsporogenesis in males and megasporogenesis in females (Alvarez-Buylla et al. 2010; Ashapkin et al. 2019). In *Arabidopsis*, the H3K4me1 is mediated by Trithorax/Thrithorax-related (Trx/Trr) and Set1-type histone methyltransferases. Most notably, ATX1 and ATX2 are known to be involved in H3K4 methylation, and ATX7 in the overall maintenance of H3K4me2/me3 and H3K36me2 (Pontvianne et al. 2010). The up-regulation of these three genes in male *S. latifolia* floral tissue (Bačovský et al. 2022) supports our observations and suggests transition into pre-meiotic phase in males (Fig. 4). In particular, the enrichment of H3K4me1 maintained by ATX1 may be indicative of a developmental switch toward meiotic competence.

H3K9me2 plays a significant role in the maintenance of non-CG DNA methylation, acting through KRYPTONITE histone-methyltransferase and chromatin remodeling DDM1 (Johnson et al. 2007; Zemach et al. 2013). Chromomethylases CMT2 and CMT3, in turn, bind H3K9me2 and use it as a signal for DNA methylation, forming a feedback loop helping to maintain inactive chromatin (Gehring and Henikoff 2008). Despite an overall decline, particularly in males, H3K9me2 remains consistently detectable across all developmental stages. We suggest that global levels of H3K9me2 are required for the efficient suppression of pericentromeric chromatin and TE elements, which are especially abundant in *S. latifolia* genome (Akagi et al. 2025; Moraga et al. 2025).

Our results of the distinct chromatin signatures, marked by the distribution of H3K4me1 and H3K9me2, along with active Pol-IIS2ph, indicate tissue-specific developmental programs associated with microspore and egg cell production. The observed cell identities within the floral meristem layers are shaped by transcriptional programs tightly coupled to chromatin state. Thus, our findings imply that gene accessibility to RNA polymerase II is modulated by histone modifications, where active (H3K4me1) and repressive (H3K9me2) marks co-occur or dynamically interact during key developmental transitions (Baroux and Autran 2015). Finally, we demonstrate that high-content imaging, besides recently adopted semi-automated image segmentation in *Arabidopsis* (Randall et al. 2022; Rutowicz et al. 2024) may allow high-throughput screening of various genetic and epigenetic traits in non-model plant species. Accurate segmentation proved essential for quantifying chromatin states in complex meristematic tissues. Standard thresholding systematically underestimated nuclei in dense regions, whereas TruAI-assisted instance segmentation reliably resolved overlapping objects and increased detection rates 2–3-fold. This methodological advance enabled high-resolution, stage-specific analysis of epigenetic dynamics in *S. latifolia* floral development. This approach provides complex data across distinct cell lineages and tissues on elevated speed and accuracy. By applying it to *S. latifolia*, we offer a comprehensive view of meristem and floral organ development, with a focus on epigenetic dynamics in spatially resolved floral tissues and sexual dimorphism. We hope that our work encourages further research on the role of epigenetic regulation in sex-specific flower development and its potential modulation during reproductive organ differentiation.

## Author contribution

Authors TJ, VB and VH planned and designed the research. TJ, VB and HPS performed the experiments. TJ and MK analyzed the data in RStudio for the statistical analysis. TJ and VB performed the visual data analysis in TIBCO Spotfire. TJ and VB wrote the main text of the manuscript with the input of all authors. All authors approved the final version of the manuscript.

## Acknowledgement

The work was supported from the project TowArds Next GENeration Crops, reg. no. CZ.02.01.01/00/22_008/0004581 of the ERDF Programme Johannes Amos Comenius.

