## Supplementary Information for "Epigenetic reprogramming guides sexual dimorphism during floral development in *Silene latifolia*"

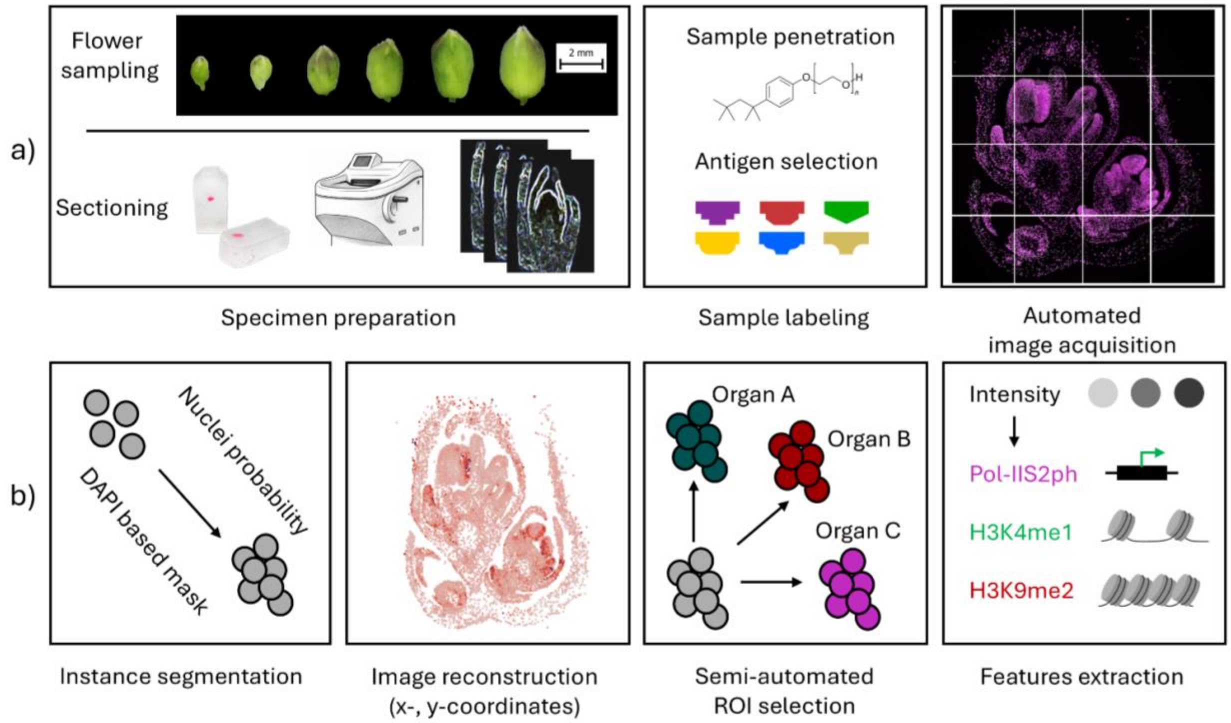


**Figure S1.** Schematic overview of specimen preparation and image analysis workflow.

(a) Dissection of male and female flower buds, followed by tissue embedding and sectioning. Specimens are examined under a light microscope and subsequently used for immunodetection and automated image acquisition. (b) Schematic representation of high-content imaging and image analysis. Images are segmented into individual objects, components, and background. Dense specimens, such as flower tissue sections, require additional post-processing through instance segmentation (see Fig. 1). Well-separated objects are then converted into x, y coordinates with corresponding maximum and mean intensity levels, allowing to assess various features (transcription activity, open or closed chromatin), which can be visualized in ScanR analysis software, Tibco Spotfire, or optionally in ImageJ/Fiji.

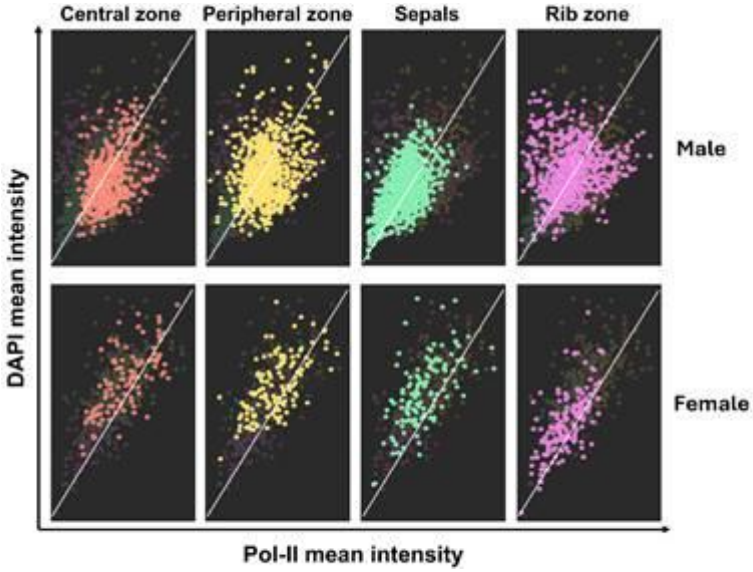


**Figure S2**. Distribution and mean intensity of Pol-IIS2ph at STG 3–4. Scatter plots showing the relationship between Pol-II mean fluorescence intensity (x-axis) and DAPI mean intensity (y-axis) across different meristematic zones in *Silene latifolia* male (top row) and female (bottom row) floral meristems. Each panel corresponds to a specific meristematic region: central zone (red), peripheral zone (yellow), sepals (cyan), and rib zone (magenta). Individual points represent single nuclei, with colors matching the respective tissue zones as assigned in TiBCO Spotfire. These plots illustrate transcriptional activity (Pol-II) in relation to nuclear content (DAPI) and highlight zone-specific differences between sexes at an early floral developmental stage. Note global M:F comparison of fluorescence intensity in Fig. 2.

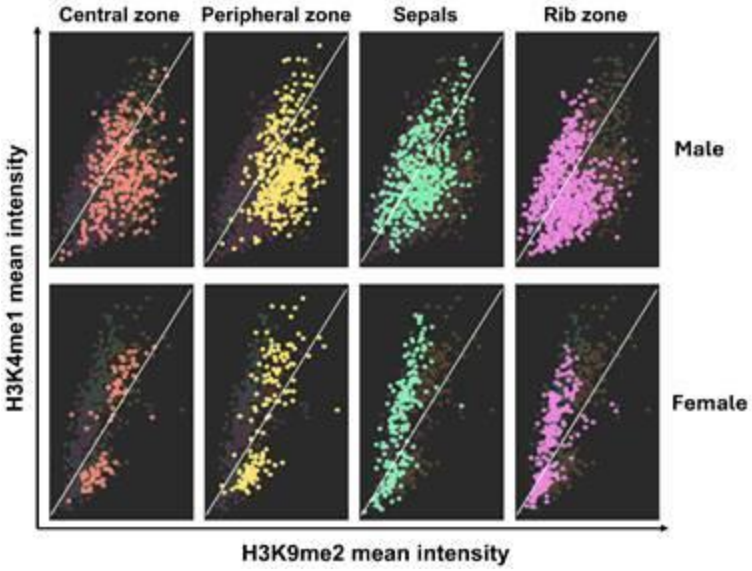

**Figure S3**. Distribution and mean intensity of H3K4me1 and H3K9me2 at STG 3–4 in male and female in *S. latifolia*. Scatter plots show the correlation between the mean intensity of the repressed chromatin-associated mark H3K9me2 (x-axis) and the active chromatin-associated mark H3K4me1 (y-axis) across four meristematic domains: central zone, peripheral zone, sepals, and rib zone. Data points represent individual nuclei quantified from male (top row) and female (bottom row) floral meristems. Colors match those used in Fig. 1. A visible inverse or distinct pattern of mark enrichment reflects zone- and sex-specific chromatin states, with females showing increased H3K4me1 in central zones and males exhibiting stronger H3K9me2 enrichment, particularly in peripheral and rib zones. Note global M:F comparison of fluorescence intensity in Fig. 2.

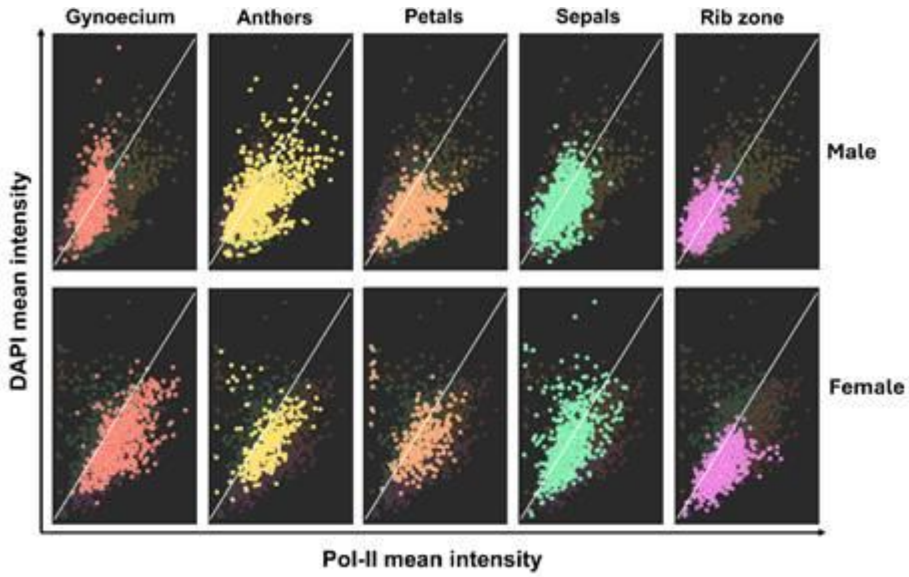


**Figure S4.** Distribution and mean intensity of Pol-II at STG 5–6. Scatter plots showing the relationship between Pol-II mean fluorescence intensity (x-axis) and DAPI mean intensity (y-axis) across different meristematic zones in *Silene latifolia* male (top row) and female (bottom row) floral organ primordia. Each panel corresponds to a specific organ: gynoecium (red), anthers (yellow), petals (orange), sepals (cyan), and rib zone (magenta). Individual points represent single nuclei, with colors matching the respective tissue zones as assigned in TiBCO Spotfire. These plots illustrate transcriptional activity (Pol-II) in relation to nuclear content (DAPI) and highlight organ-primordia specific differences between sexes. Note global M:F comparison of fluorescence intensity in Fig. 3.

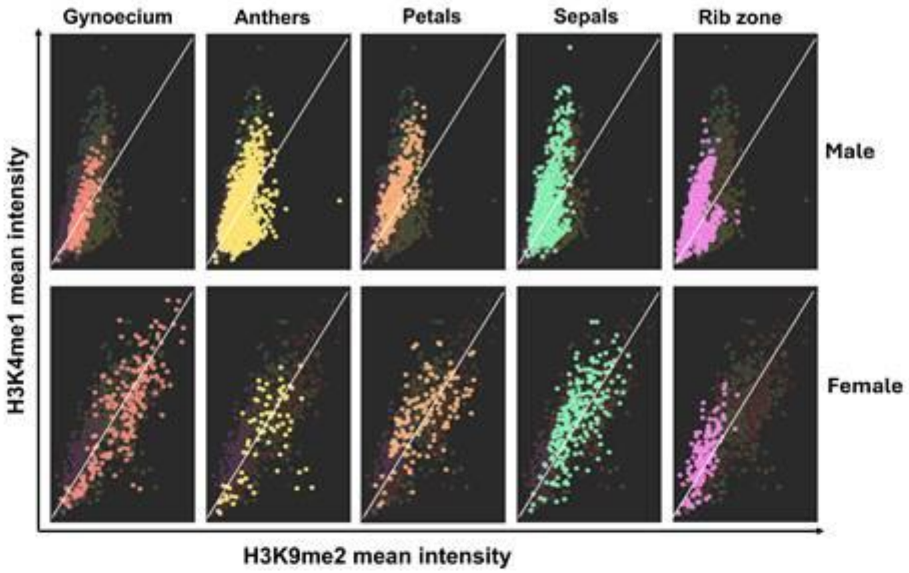


**Figure S5.** Distribution and mean intensity of H3K4me1 and H3K9me2 at STG 5–6 in male and female in *S. latifolia*. Scatter plots show the correlation between the mean intensity of the repressed chromatin-associated mark H3K9me2 (x-axis) and the active chromatin-associated mark H3K4me1 (y-axis) across specific organ primordia: gynoecium (red), anthers (yellow), petals (orange), sepals (cyan), and rib zone (magenta). Data points represent individual nuclei quantified from male (top row) and female (bottom row) floral organs. Colors match those used in Fig. 3. A visible inverse or distinct pattern of mark enrichment reflects zone- and sex-specific chromatin states, with females showing increased H3K4me1 in gynoecium and petals, and males exhibiting stronger H3K4me1 enrichment in anthers, petals and sepals, but with lower intensity compared to female organs (Fig. 3c, d).

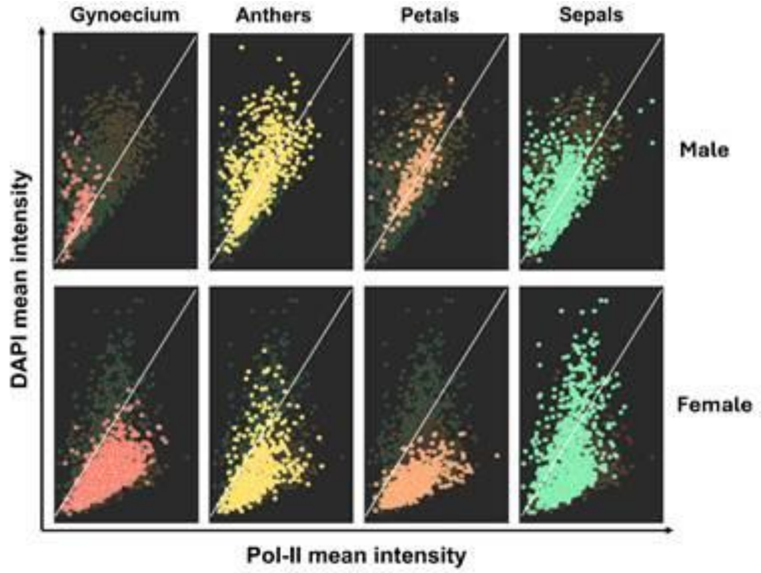

**Figure S6.** Distribution and mean intensity of Pol-II at STG 7–8. Scatter plots showing the relationship between Pol-II mean fluorescence intensity (x-axis) and DAPI mean intensity (y-axis) across different meristematic zones in *Silene latifolia* male (top row) and female (bottom row) floral organ primordia. Each panel corresponds to a specific organ: gynoecium (red), anthers (yellow), petals (orange) and sepals (cyan). Individual points represent single nuclei, with colors matching the respective tissue zones as assigned in TiBCO Spotfire. These plots illustrate transcriptional activity (Pol-II) in relation to nuclear content (DAPI) and highlight organ-primordia specific differences between sexes. Global expression increases toward anthers and sepals in male, and particularly in gynoecium and sepals in females. Note global M:F comparison of fluorescence intensity in Fig. 4.

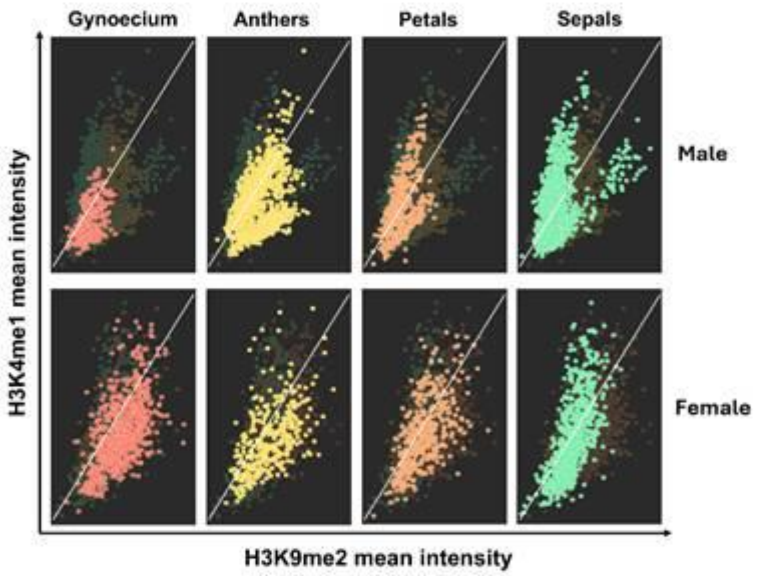


**Figure S7.** Distribution and mean intensity of H3K4me1 and H3K9me2 at STG 7–8 in male and female *S. latifolia*. Scatter plots show the correlation between the mean intensity of the repressed chromatin-associated mark H3K9me2 (x-axis) and the active chromatin-associated mark H3K4me1 (y-axis) across specific organ primordia: gynoecium (red), anthers (yellow), petals (orange) and sepals (cyan). Data points represent individual nuclei quantified from male (top row) and female (bottom row) floral organs. Colors match those used in Fig. 4. Anthers, petals and sepals display larger enrichment compared to gynoecium in males. Gynoecium in females has larger enrichment of H3K9me2. Note global M:F comparison of fluorescence intensity in Fig. 4.
